# Effects of sexual dimorphism on pollinator behaviour in a dioecious species

**DOI:** 10.1101/2021.04.15.440026

**Authors:** L. Moquet, A-L Jacquemart, M. Dufay, I. De Cauwer

**Affiliations:** Univ. Lille, CNRS, UMR 8198 - Evo-Eco-Paleo, F-59000 Lille, France; UMR PVBMT, CIRAD, Saint Pierre, La Réunion, France; Genetics, Reproduction, and Populations research group, Earth and Life Institute, Université catholique de Louvain, B-1348, Louvain-la-Neuve, Belgium; CEFE, Univ Montpellier, CNRS, EPHE, IRD, Montpellier, France

**Keywords:** Dioecy, floral trait sexual dimorphism, pollination, visitation sequence, pollen transfer, *Silene dioica*

## Abstract

Floral traits often display sexual dimorphism in insect-pollinated dioecious plant species, with male individuals typically being showier than females. While this strategy is theorized to be optimal when pollinators are abundant, it might represent a risk when they become scarce, because the disproportionately high number of visits on the most attractive sex, males, might preclude efficient pollen transfer from males to females. Here, the effect of sexual dimorphism on pollination efficiency was assessed in experimental arrays of dioecious *Silene dioica* that were exposed to one frequent visitor of the species, *Bombus terrestris*, and that differed in the magnitude of sexual dimorphism for either flower number or flower size. While flower size dimorphism did not impact pollination efficiency, we found that flower number dimorphism negatively affected the number of visits on female plants, on female flowers and on the number of female flowers visited after a male flower. However, flower number dimorphism had no effect on the number of pollen grains deposited per stigma, presumably because the decrease in the number of visits to female flowers was compensated by a higher number of pollen grains deposited per visit.

## Introduction

One common feature in plants with separate sexes is sexual dimorphism, both in terms of vegetative and reproductive traits (reviewed in Barrett and Hough 2013). In insect-pollinated dioecious species, male plants often display more conspicuous floral phenotypes than females, at least in temperate species, with males typically producing larger floral displays and/or larger flowers (Guitián 1995, Delph et al. 1996, Pailler et al. 1998, Eckhart 1999, Costich and Meagher 2001, Ramsey and Vaughton 2001, Kriebel 2014, Matsuhisa and Ushimaru 2015, Cuevas et al. 2017). One common explanation for this pattern is sexual selection: if male’s reproductive success is more limited by mate acquisition than female’s reproductive success (as predicted by Bateman principles, Bateman 1948), any trait involved in pollinator attraction should be under stronger selection in males than in females (Delph and Ashman 2006, Moore and Pannell 2011).

Pollinators have been shown to preferentially visit male plants in several dioecious species (e.g. Ågren et al. 1986, Carlsson-Granér et al. 1998). This preference can be driven by the difference between males and females in terms of flower number or size (e.g. Vaughton and Ramsey 1998, Bond and Maze 1999, Pickering 2001) and/or associated with other male specificities, such as the occurrence of stamen and pollen (Lunau 2000, Nicholls and Hempel de Ibarra 2017). One potential consequence of unbalanced visitation rates between sexes is that females might, on average, not receive enough visits for optimal pollination. When female’s reproductive success is pollen limited, the strength of selection on attractive floral traits in females is expected to increase (Sletvold and Ågren 2016, Caruso et al. 2019), which could potentially result in a decrease in the magnitude of sexual dimorphism. However, an abrupt decrease in pollinator abundance could have deleterious demographic effects well before any evolutionary response could take place. Accordingly, the modelling study by Vamosi and Otto (2002) predicts that pollinator preference towards one sex, in this case males, could lead to population extinction when pollinators are scarce and when plants display a high degree of sexual dimorphism.

To date, only a few experimental studies have examined the effect of sexual dimorphism on pollinator behaviour at the plant population level. By manipulating the distribution of floral size between males and females in experimental arrays of the dioecious *Sagittaria latifolia*, Glaettli and Barrett (2008) showed that the proportion of pollinator visits on males increased with the magnitude of floral size sexual dimorphism. This pattern resulted from an increase in the number of visits on males, with no change in the number of visits on females. Based on their results, the authors argued that males might benefit from showier flowers in terms of pollen export, whereas increased sexual dimorphism should not impact female reproductive success in this system. In a similar study, Vaughton and Ramsey (1998) manipulated the occurrence of flower number dimorphism in *Wurmbea dioica* and found that pollinators preferred males even when flower number was the same in both sexes, possibly due to the larger flower sizes in males. However, since no pollen limitation was detected, this preference for males should not have incurred any demographic cost in the study population. As underlined by these two studies, the effect of sexual dimorphism on the number of visits received by females is not necessarily a good predictor of pollination efficiency. To test the hypothesis that the magnitude of sexual dimorphism negatively impacts pollination, one also needs to measure pollen transfer, which depends not only on the number of visits on females, but also on the quantity of pollen carried by insects, which might in turn depend on whether or not insects visited male flowers before reaching female flowers. Yet, the order of visits on male and female flowers by pollinators has rarely been investigated (but see Tsuji et al. 2020), and the consequences of sexual dimorphism on the ability of pollinators to efficiently transfer pollen from male to female flowers remains unclear and largely untested.

In the current study, we examined the effect of sexual dimorphism on pollinator behaviour and pollen transfer in *Silene dioica*. This species shows sexual dimorphism for a variety of traits, including floral size and floral number, with males producing markedly larger and more abundant flowers than females (Kay et al. 1984, Moquet et al. 2020). In this species, males have been shown to attract more pollinators than females, possibly as a consequence of their showier displays (Kay et al. 1984, Carlsson-Granér et al. 1998). Here, we independently manipulated the magnitude of sexual dimorphism in flower number and flower size in experimental populations and quantified the effect of this variation on the visitation patterns of *Bombus terrestris*, a frequent visitor of the study species *in natura* (Kay et al. 1984). Because the negative effects of sexual dimorphism on pollination efficiency are expected to occur when pollinators are scarce (Vamosi and Otto 2002), the number of pollinator visits was fixed and corresponded to the number of flowers in each experimental plant population. By comparing arrays where males and females showed no sexual dimorphism with arrays in which sex dimorphism occurred for only one of the two traits, we were able to disentangle the effects of flower number and flower size on visitation patterns. We measured the number of visits on male and female plants and flowers, the visiting sequences (order of visits on male and female flowers), and the pollen transfer to stigmas. Our prediction is that pollination efficiency will be lower in sexually dimorphic experimental populations than in populations where there are no differences between males and females in terms of floral visual signalling.

## Material and methods

### Experimental plants and bumblebee species

*Silene dioica* is a common perennial herb of north-western Europe, flowering from April to June (Kay et al. 1984). This species has a generalist insect-pollination system, with *Bombus* species and syrphids as most common visitors (Baker 1947). Female flowers have five stigmas and male flowers ten stamens. Males typically carry showier displays than females: on average, they produce over ten times more flowers over the course of the flowering season, with corollas that are 20% wider than in females (Moquet et al. 2020). Both male and female flowers secrete nectar at the base of the corolla (Vogel 1998, Comba et al. 1999), with male flowers producing less abundant but more concentrated nectar than female flowers (Kay et al. 1984, Comba et al. 1999). We used the same experimental material as in Moquet *et al*. (2020): plants originated from seeds collected in July 2015 and 2016 in eight natural populations in northern France. Seeds were sown in autumn the same year they were harvested. Prior to the experiment, which was carried out in spring 2017, plants were vernalized for 10 weeks at 6 °C (Lille University, France) and placed in an experimental garden until the flowering season (catholic University of Leuven, Belgium).

*Bombus terrestris* is a frequent visitor of *S. dioica* (Baker 1947, Kay et al. 1984, Goulson and Jerrim 1997). We used colonies of *B. terrestris* from BioBest Biological Systems (Westerlo, Belgium) in which we removed access to the factory-supplied nectar reservoir during the experiment to encourage foraging. Pollen was delivered *ad-libidum* directly to the nests.

### Experimental design

To test whether the occurrence of sexual dimorphism in flower number and/or size impacted visitation rates, pollinator behavior, and pollen transfer from male flowers to female stigmas, we ran experiments in an indoor flight arena (2.80 x 2.20 x 2.00 m, L x W x H) in May and June 2017. For each experimental session, we selected ten plants, five females and five males, according to their characteristics in terms of corolla diameter and flower number on the observation day. We counted the number of flowers and measured corolla diameter on five randomly selected flowers with a digital calliper precise to 0.01 mm.

When less than five flowers were available, all the flowers were measured. The 24 observation sessions were distributed across three treatments (Figure 1):

i. male and female corolla of equivalent size, more male flowers than female flowers (flower number dimorphism, n=8)
ii. male corolla larger than female corolla, an equal number of flowers (flower size dimorphism, n=8 sessions), and
iii. male and female corolla of equivalent size, an equal number of flowers (no dimorphism, n=8 sessions).

**Figure 1 :**
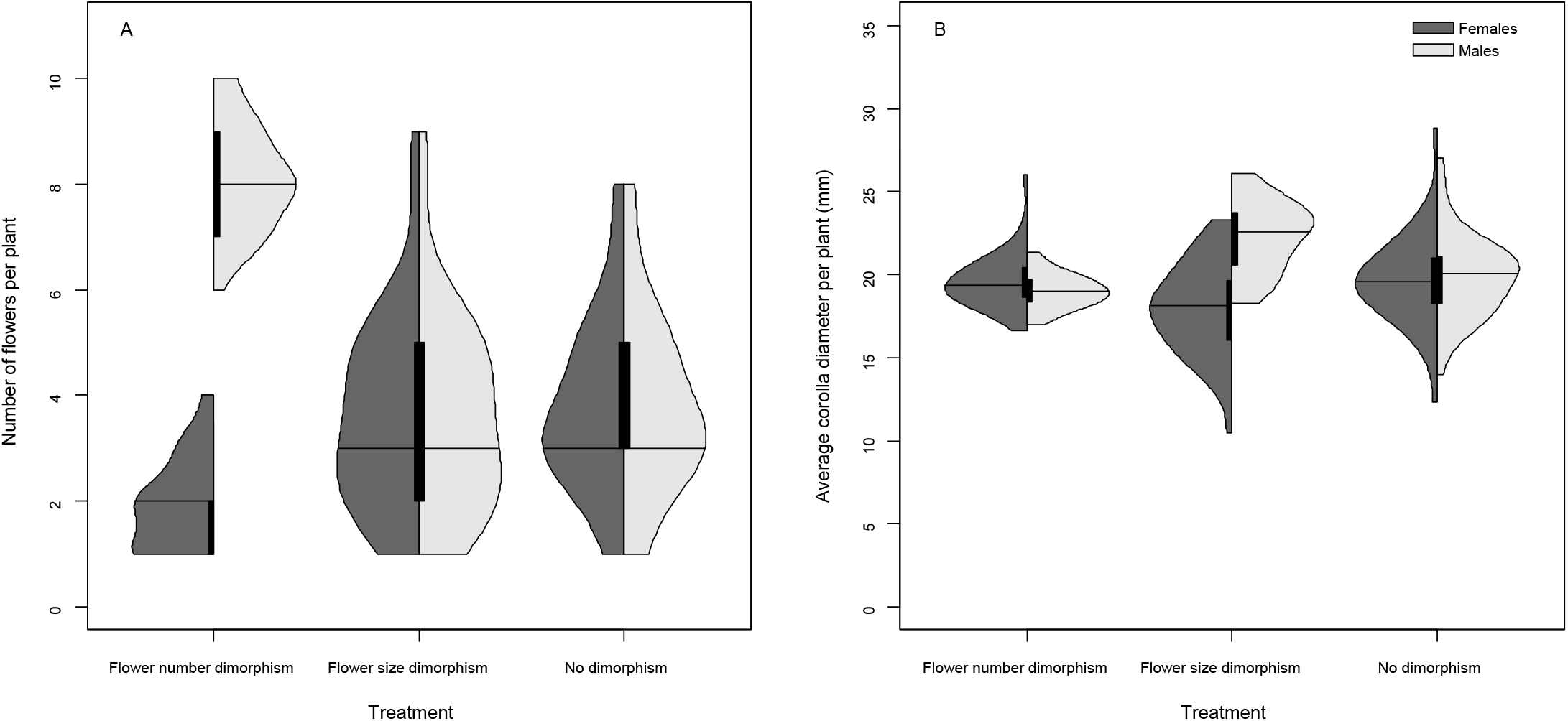
Violin plots showing the distribution of (A) flower number and (B) corolla size in male and female plants in the three treatments.

Within the flight arena, plants were arranged in four rows (row 1 and 3: three plants, row 2 and 4: two plants). Within and between rows, neighboring plants were separated by exactly 40 cm. Male and female plant positions were randomly arranged each session.

The day before the experiment, plants were placed in a greenhouse to exclude nectar-feeding insects and allow nectar replenishment. Flower buds were marked with a water-based marker pen in order to identify the virgin flowers that were used to estimate pollen loads after each observation session.

### Training phase

Three different hives were used, alternating hive × treatment combinations. Before the experiment, bumblebees were allowed to familiarize with the flight arenas and with reward collection. During this training phase, one hive was open, and bumblebees could move freely. Bumblebees were trained on six plants (3 males, 3 females) randomly arranged in the flight arena. In order to limit the establishment of a plant-sex preference, plants were selected to have similar characteristics (plant size, flower numbers, and corolla size). We ran five training sessions of 30 min for each colony before the observation sessions, then one 30 min training session every two weeks.

### Observation protocol

Each observation session, five individual bumblebees were released one by one, in the flight arena. One observer, posted behind a net, noted the bumblebees visiting sequences. Any occurrence of a bumblebees probing a flower was recorded as a visit. Bumblebees were captured and marked after they visited ten flowers. In the rare cases where bumblebees did visit fewer than ten flowers and stopped foraging for more than 10 minutes, the trial was stopped and a sixth trial was performed. Each session consisted of exactly 50 visits (five bumblebees × ten visits) in the flight cage. Because the maximum number of flowers per session was 50 for all treatments, all flowers had a chance to be visited at least once. Some flowers were visited more than once, thus, the amount of reward of an individual flower could decline during each session. This experiment thus mimics common situations in which nectar secretion rates are weak in comparison to visitation rates (Real 1981, Ashworth and Galetto 2002, Barp et al. 2011).

### Pollen transfer

One day after the experiment, we individually collected styles of one or two marked flowers per female plant in order to analyse the quantity of pollen deposited onto the stigmas. Styles were kept in FAA (ethanol 70%: formaldehyde 35%: acetic acid; 8:1:1) until analysis. Before observation, styles were rinsed with demineralised water for one hour, soaked in a sodium hydroxide (4M) for three hours, and again rinsed in water for approximatively one hour. Styles were then put on a microscope slide with a drop of aniline blue solution (0.87 g KH2PO4, 0.1 g of aniline blue, 50 ml water) and pollen grains were counted under a fluorescence microscope (excitation filter 420–440 nm, emission filter 480 nm, Nikon Eclipse).

### Statistical analysis

Generalized linear mixed models (GLMM) were used to test for the effects of sexual dimorphism on 1) the distribution of visits between male and female plants during a session, 2) the characteristics of the visit sequence performed by each bumblebee and 3) the pollen transfer from male to female flowers. For all following analyses, the effects of “flower number dimorphism” and “flower size dimorphism” treatments were assessed separately using comparisons with the “no dimorphism” treatment. For each model, hive and observation session were primarily added as random factors and were removed when they had no influence on our models. All models were realised with the lme4 package in R version 4.0.2 (R Development Core Team 2008).

#### Distribution of visits among male and female plants

We tested whether plant sex, treatment and sex × treatment interaction had an influence on the number of bumblebee visits received by each plant. In these models, the individual was the unit of analysis (n = 80 plants per treatment). Two response variables were considered: i) the total number of visits per plant and ii) an estimate of the number of visits per flower (number of visits on one plant divided by the number of open flowers on this plant) during one session. We used a negative binomial distribution for the first model and a Gaussian distribution for the second one.

#### Sequences of visits

We tested for the effect of treatment on several characteristics of the visit sequence performed by each bumblebee. In these models, bumblebee foraging bouts were thus the unit of analysis. Only complete sequences (i.e. 10 flowers visited) were kept (n = 34, 36 and 39 sequences for “flower number dimorphism”, “flower size dimorphism” and “no dimorphism” treatments, respectively). The response variables were: (i) the number of visited females, (ii) the number of visited female flowers, (iii) the number of visited female flowers divided by the total number of female flowers available in the experimental plot, (iv) the number of female flowers visited after the insect had visited at least one male flower, which provides an estimate of the number of potentially pollinating visits, and (v) the sex of the first visited flower. Moreover, because there was no time for nectar replenishment between bumblebee trials, we added bumblebee flying order (from 1 to 5) as an explanatory variable in the model, as well as the treatment × order interaction. We used binomial distributions for models analyzing variable (v), gaussian for models analysing variable (iii) and Poisson distributions for the other variables.

Regarding the number of female flowers visited after at least one male flower, this variable should depend on both the number of female flowers in the sequence and the order of the visits. For each observed visit sequence, we thus simulated 1000 random sequences of 10 flowers and containing the same number of female flowers. We then measured the proportion of simulated sequences that had a lower number of visited females after at least one male compared to the observed sequence. When this proportion was lower than 0.05, we considered the sequence to depart from expectations under random behaviour of the insect.

#### Pollen deposition

To test whether floral sexual dimorphism affected pollination success, we analyzed the effects of the treatment on the number of pollen grains deposited on collected stigmas. When more than one stigma was collected, pollen quantity was averaged across stigmas (n = 40 females for the “flower number” treatment, n = 36 for the two other treatments). Because a low number of open flowers on a given female potentially concentrates all deposited pollen grains on the same stigmas, we added the number of open flowers as a co-variable and tested for the interaction between treatment and flower number. We used a distribution of Poisson in these models.

## Results

### Does sexual dimorphism affect the distribution of visits on male and female plants?

When comparing the number of visits received per plant between the “flower number dimorphism” and the “no dimorphism” treatment, we detected an effect of both plant sex (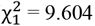, *P* = 0.002) and the interaction between plant sex and treatment (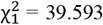, *P* < 0.001). In the sessions with no sexual dimorphism, female plants received slightly more visits than male plants, and this pattern was reversed when male plants carried more flowers (Fig. 2A). Only plant sex had a significant effect (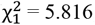, *P* = 0.016) on the number of visits per flower, with higher visitation rates on females in both treatments (Fig. 2B).

**Figure 2 :**
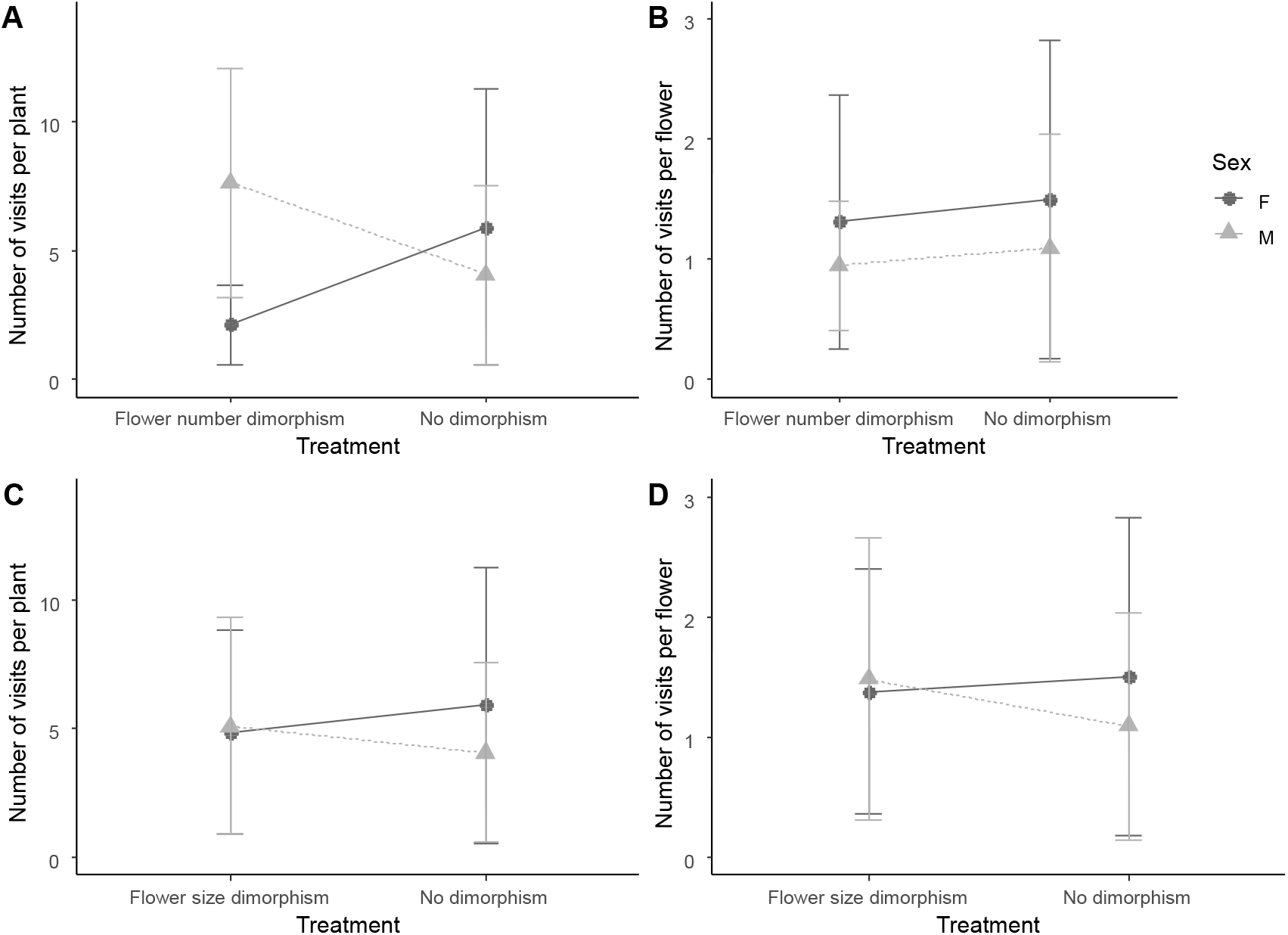
Number of visits received per plant and per flower, for females (dark grey dots) and for males (light grey triangles). A and B show comparisons between the “no dimorphism” and the “flower number dimorphism” treatment for the number of visits at the plant and flower level respectively. C and D show comparisons between the “no dimorphism” and the “flower size dimorphism” treatment for the number of visits at the plant and flower level respectively.

Regarding the comparison between the “flower size dimorphism” and the “no dimorphism” treatments, although the number of visits per plant and per flower was similar in males and females when males produced larger flowers, the interaction between plant sex and treatment was not significant (Fig. 2C and 2D).

### Does sexual dimorphism impact the characteristics of visitation sequences?

Bumblebee behavior was found to be statistically similar between the “flower size dimorphism” and the “no dimorphism” treatments for all tested variables. On the contrary, when comparing bumblebee behavior between the “flower number dimorphism” and the “no dimorphism” treatment, we found significant differences for several metrics (Fig. 3). Sexual dimorphism for flower number reduced the number of visited female plants (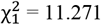, *P* < 0.001) and female flowers (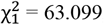, *P* < 0.001), as well as the number of female flowers visited after a male flower (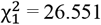, *P* < 0.001). The sex of the first visited flower was also affected (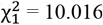, *P* = 0.002), with a first visit on a male flower in 74% of sequences in the “flower number dimorphism” treatment, against 36% of sequences in the “no dimorphism” treatment. Treatment did not affect the number of visited female flowers when corrected by the total number of female flowers available in the arena (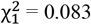, *P* = 0.068). Bumblebee flying order and the order × treatment interaction did not impact any of the tested response variables (*P* > 0.05).

**Figure 3:**
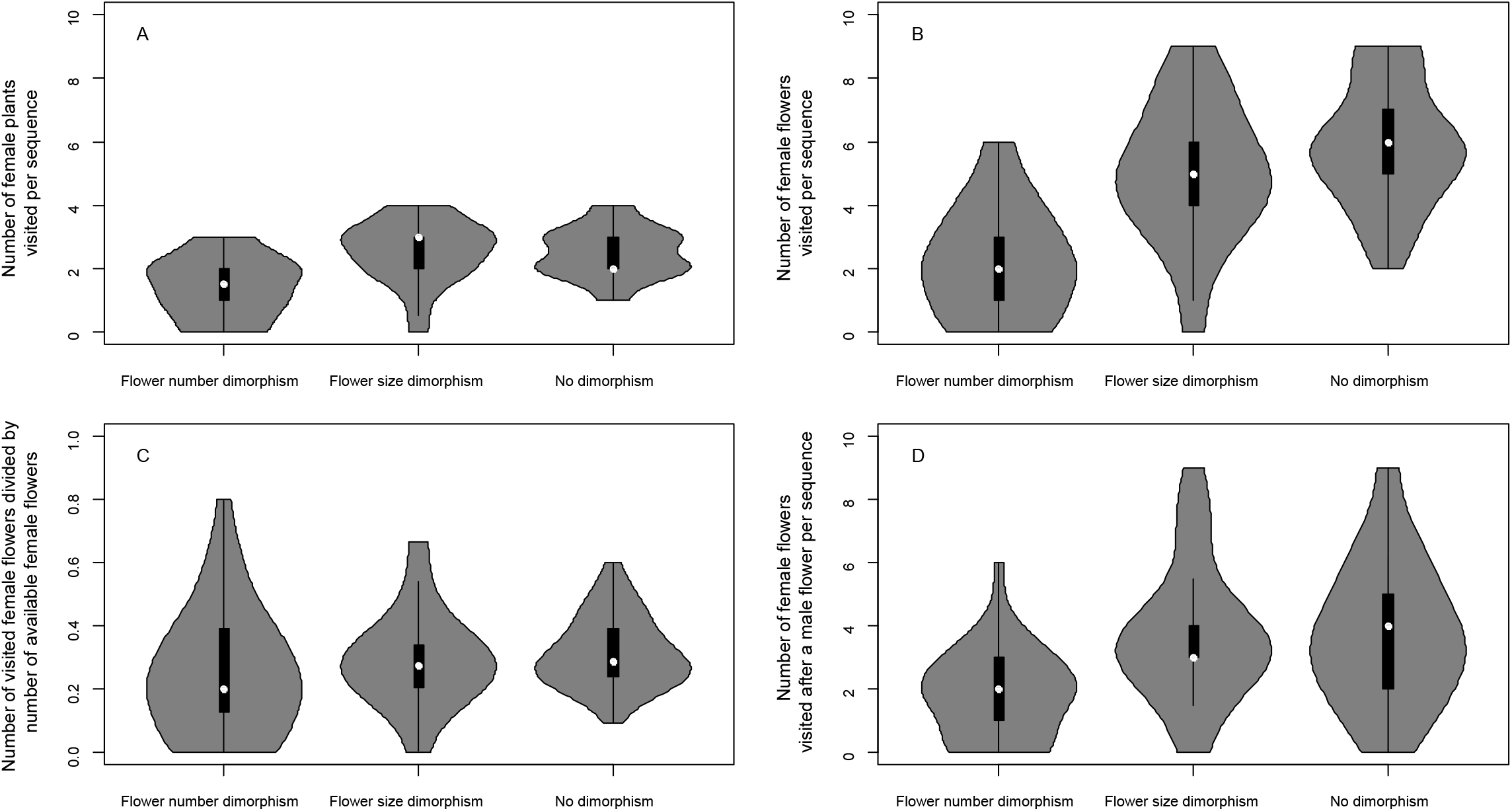
Violin plots showing the variation in visitation sequences among treatments, with the distribution of: (A) the number of female plants visited per insect (B) : the number of female flowers visited per insect, (C) : the number of visited females flowers divided by the total number of female flowers available in the arena and (D) : the number of female flowers visited after one or several male flower(s).

The number of female flowers visited after at least one male flower was significantly lower than expected under random behaviour in a fifth of the visitation sequences in the “no dimorphism” treatment (8 of the 39 observed sequences). In these sequences, visits started by one or several female flower(s), thus reducing the number of potentially pollinating visits: overall, in this treatment, 87 of the 230 visited females’ flowers were visited before any visit on a male flower, implying that 38% of visited female flowers could not receive any pollen. Such deviations from random expectations were observed only twice in the “flower size dimorphism” treatment, and never in the “flower number dimorphism” treatment.

### Effects of floral dimorphism on pollination success

Both in case of flower number and flower size dimorphism, treatment, number of flowers and the interaction between these two variables had no influence on the average number of pollen grains deposited on collected stigmas (*P* > 0.05, Fig. 4).

**Figure 4:**
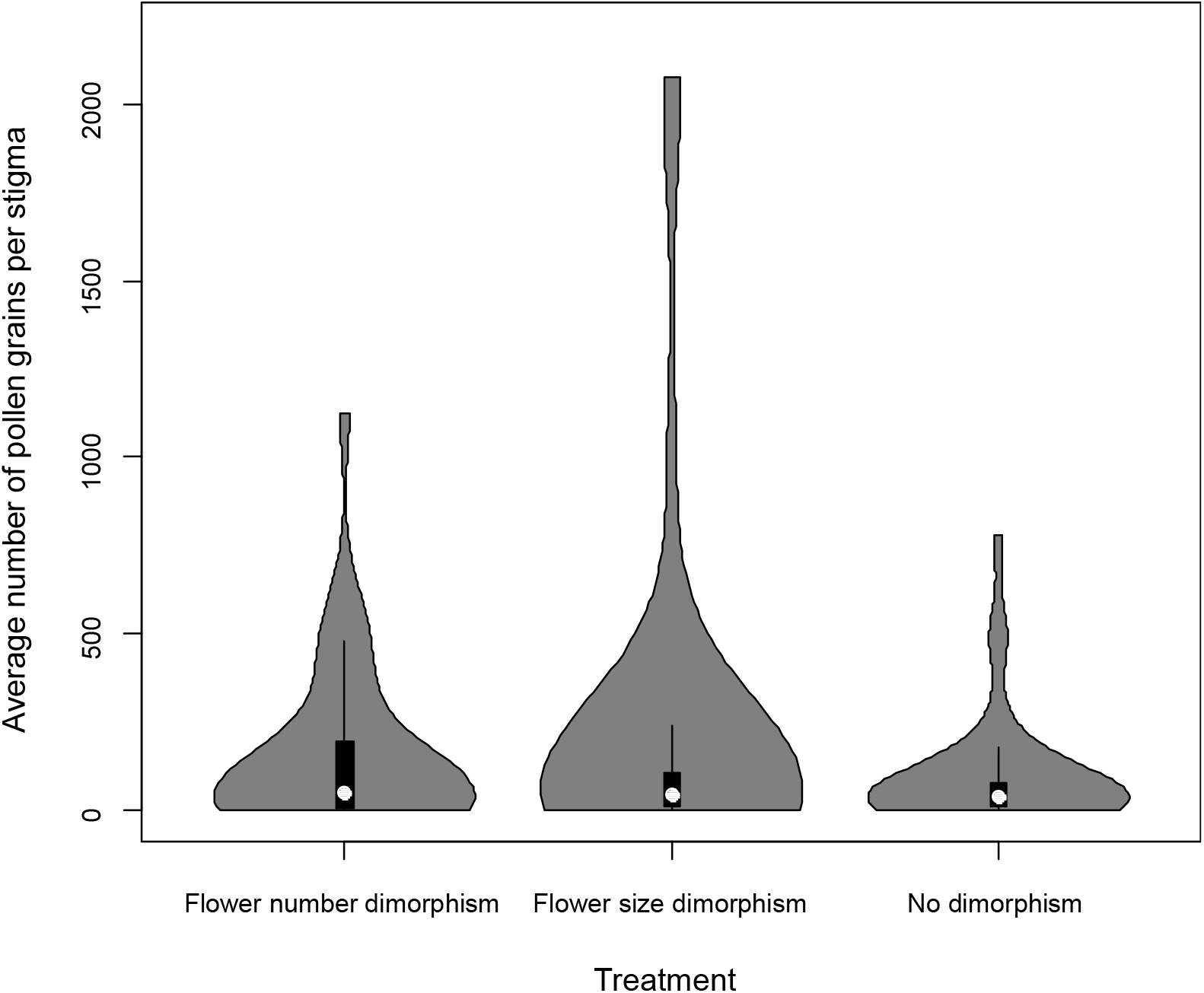
Violin plots showing the distribution of the number of pollen grains per stigma in each treatment. When more than one stigma was collected, pollen counts were averaged across stigmas.

## Discussion

While the effects of floral traits on pollinator attraction have been studied extensively (e.g. Conner and Rush 1996, Spaethe et al. 2001, Martin 2004, Hegland and Totland 2005, Ishii 2006), the response of pollinators to between sexes variation in floral phenotypes is less well understood, despite the fact that sexual dimorphism for floral traits is a common feature in insect-pollinated dioecious plants (e.g. Delph et al. 1996, Eckhart 1999). In *S. dioica*, as in many dioecious plant species, males are typically showier than females, with larger floral displays both in terms of flower number and flower size (Kay et al. 1984, Moquet et al. 2020), and flower visitors of *Silene dioica* have been shown to prefer male over female plants under natural conditions (Kay et al. 1984, Carlsson-Granér et al. 1998). In our study, by separately examining cases where sexual dimorphism occurs only for flower number or for flower size, and by comparing the visitation patterns in these two situations with trials with no sexual dimorphism, we were able to disentangle the effects flower number vs. flower size sexual dimorphism.

Two important features of our experimental design should be kept in mind when interpreting the results. First, although *S. dioica* is known to display a generalist insect-pollination system (Baker 1947, Kay et al. 1984), only one pollinator species was used in the experiment. *Bombus terrestris* was selected because the existing literature describes this species as a common visitor of *S. dioica* (Kay et al. 1984), but different pollinator taxa can display different preferences in terms of floral signals (e.g. Glaettli and Barrett 2008, Gómez et al. 2008), which could in turn impact their response to the level of sexual dimorphism. Second, because the experiments were conducted in a flight arena with fixed numbers of pollinators and visits per pollinator, we could not capture the effects of sexual dimorphism on the overall number of visits at the plant patch level. This overall number of visits should indeed depend both on the number of pollinators visiting the patch (which can increase if males are more attractive, as found by Glaettli and Barrett 2008) and on the length of visitation sequences. Keeping this in mind, we were able to detect interesting effects of sexual dimorphism on pollinator behaviour.

### Experimentally cancelling sexual dimorphism for flower number and size allowed uncovering an innate preference for female flowers

In the “no dimorphism” treatment, we observed an innate preference of bumblebees for female flowers, both when considering visits at the plant and at the flower level. Pollinators preference for female flowers in gender dimorphic species is rare although it has been observed in a few cases (e.g. gynodioecious *Fuschia thymifolia*, Cervantes *et al*. 2018; dioecious *Catasetum arietinum*: Brandt *et al*. 2020). The preference towards *S. dioica* females might be driven by differences in traits that were not measured in the current study, such as nectar production or scent emission. In this species, female flowers produce more abundant but less concentrated nectar than males (Kay et al. 1984). The quantity of sugar per flower was measured in plants from the same collection we used in this experiment in another study (Barbot et al. submitted) and it was observed that, at the flower level, reward is higher in females than in males, which might drive the preference for female flowers. Regarding olfactory signals, while scent emission and composition have been characterized in *S. dioica* (Jürgens 2004, Page et al. 2014), between sex variation remains unknown to date. In the sister species *S. latifolia*, male flowers release larger amounts of scent than female flowers, with a marked dimorphism for pollinator-attracting compounds (Waelti et al. 2009), a pattern that is common in dioecious insect-pollinated plants (Ashman 2009). Further studies will be needed to determine whether or not scent is sexually dimorphic and whether this trait plays a role in pollinator attraction in *S. dioica*.

The preference for females in the “no dimorphism” treatment resulted in a higher number of sequences starting by visits on one or several female flowers. This was mainly due to a probabilistic effect: with more female flowers in the sequence, the probability of starting by a female flower increases. In addition, we found that some sequences featured less females visited after a male than expected under random behaviour, which could be driven by a slight active preference for female flowers. As a result, for 38% of visited female flowers in the « no dimorphism » treatment, the pollination sequence did not feature an upstream visit to a male flower, highlighting the fact that a higher attractivity of females might negatively impact pollination efficiency.

### Flower size dimorphism had no effect on pollinator visitation patterns

Flower size has been shown to play a role in pollinator attraction in many other systems (e.g. Bell 1985, Galen and Newport 1987, Stanton and Preston 1988). This trait can indeed directly impact the intensity of the visual signal and/or the quantity of reward for pollinators in systems where flower size is associated with reward abundance (Blarer et al. 2002, Brunet et al. 2015). Since no significant preference for males was found when they carried larger flowers, the effect of this trait on pollinator attraction was not supported in our study. One possible explanation is that, if sugar quantity was on average higher in female flowers as expected based on other studies (Barbot et al. submitted), this may have prevented the emergence of a clear preference for large flower sizes. Indeed, we observed a tendency for sexual dimorphism in flower size to offset the innate preference of pollinators for females, but this effect was not significant.

Another study on the same collection of plants showed positive directional selection acting on flower size in males in open pollination conditions (i.e. siring success increased with flower size), although selection intensity appeared weak compared to other traits (Barbot et al. submitted). The apparent discrepancy between our results and the positive selection gradient on flower size detected in males might originate from the fact that our experimental design relied on a minimization of within-sex phenotypic variance in each treatment, implying that the showiest male phenotypes were excluded here.

### Flower number dimorphism affected pollinator preference and some features of the pollination sequences, but not pollen transfer

Flower number sexual dimorphism is probably the most notable dimorphism in *S. dioica*. Its magnitude strongly varies across the flowering season, but it is always male biased with a floral sex-ratio that can reach 1:30 at the end of the flowering season (Moquet et al. 2020). In our experiment, males carried on average four times more flowers than females, which is a relatively low bias in floral sex-ratio compared to what pollinators might encounter in natural populations. In our study, flower number dimorphism cancelled the innate preference of pollinators for females and affected several metrics of bumblebee visitation sequence.

Compared to the “no dimorphism” treatment, male plants in the “flower number dimorphism” treatment received significantly more visits than females, suggesting that the documented preference of pollinators for male plants in this species might mainly be driven by their larger floral displays. Pollinator preference for individuals with large number of flowers makes sense in the light of optimal foraging theory: pollinators should prefer plants with higher flower density because the energy and time to forage on them are lower than on plants with sparser flowers (Thomson 1988, Campbell 1989). Male biased preference was not detected at the flower level, which was probably due to a decelerating relation between the number of flowers and the number of visits (Klinkhamer and de Jong 1990), as found in many other plant species (Robertson 1992, Conner and Rush 1996, Vaughton and Ramsey 1998). At the plant level, whether or not this male-biased preference in settings where males have larger number of flowers benefits siring success should depend on the quantity of pollen removed by insects during each visit on a single flower, as well as on the overall abundance of pollinators visiting the population (de Jong and Peter 1994). The fact that flower number did not impact siring success in *S. dioica* (Barbot et al. submitted) suggests that the higher number of visits experienced by showy males might not necessarily translate into a higher siring success (de Jong and Peter 1994).

Interestingly, when we standardized the number of visited female flowers with the total number of female flowers available in each session, we did not record any effects of sexual dimorphism. These results suggest that the observed pattern is not due to any avoidance of female flowers by bumblebees or active preference for male flowers, but mainly to a decrease in the probability of encountering a female flower in the arena when the number of male flower increases.

How did this translate in terms of pollen transfer from male to female flowers? Our observations suggest that flower number sexual dimorphism can have conflicting effects on pollination efficiency. Because pollinators visited male flowers more often when male plants had more flowers than females and started their sequence by a male flower much more frequently, the proportion of visited female flowers potentially receiving pollen (i.e. visited after one or several male flowers) was higher with sexual dimorphism. However, because the overall number of visited female flowers was much lower, we found the absolute number of potentially pollinated females to be negatively impacted by sexual dimorphism.

This result is consistent with the theoretical predictions made by Vamosi and Otto (2002), who argued that decreasing the relative attractivity of female plants should diminish the number of pollinating visits. One other effect of this higher number of visited male flowers could be that pollinators accumulate a high number of pollen grains on their body. Pollen load on insects should depend on various attributes of both pollen grains and insect morphology and behaviour, which was not be measured during the current study. Pollen carryover is generally considered to be high in flowering plants (Robertson 1992). This could explain why no difference was found in the average number of pollen grains deposited on stigmas among treatments. We can suggest that, while the number of potentially pollinating visits decreased with sexual dimorphism, each visit might have been more efficient in terms of pollen deposit. Such quantitative effect was not considered by the model of Vamosi and Otto (2002), which assumed a direct link between the number of pollinating visits and population viability. Other studies found no clear parallel between the number of pollinators visits and pollen receipt of stigmas. For example, in monoecious *Sagittaria trifolia*, Huang *et al*. (2006) showed that female flower attractivity to pollinators depended on floral display size and was lower than male plants, but with no clear effect on pollen receipt. The authors explained this result by stigmas being saturated with pollen even after a limited number of pollinators visits. This was not the case in the current study, as evidenced by the high number of stigmas having no or only few pollen grains. In our case, the number of male flowers visited by insects before reaching a female flower was probably critical to explain the variance in the number of pollen grains deposited on stigmas, and may explain the absence of difference between treatments.

To our knowledge, this is the first study to manipulate the occurrence of sexual dimorphism in floral traits and to study its influence on both insect behaviour and pollen transfer. Our results show that analyses of visit sequences and pollen transfer are necessary to understand the possible consequences of sexual dimorphism on pollination success. A higher number of visits on the most attractive sex did not automatically reduce pollen transfer, highlighting the fact that modelling or measuring pollinators visits may not be sufficient to predict the effect on plant reproduction.

## Acknowledgments

We are grateful to Eric Schmitt and Fanny Raux for their help in collecting the data, as well as to the “Plateforme serre, cultures et terrains expérimentaux - Université de Lille” for the support in plant collection maintenance. This work is a contribution to the CPER research project CLIMIBIO that funded L.M.’s salary. The authors thank the French Ministère de l’Enseignement Supérieur et de la Recherche, the Hauts de France Region and the European Funds for Regional Economical Development for their financial support to this project.

